# A glance at recombination hotspots in the domestic cat

**DOI:** 10.1101/028043

**Authors:** Hasan Alhaddad, Chi Zhang, Bruce Rannala, Leslie A. Lyons

**Affiliations:** College of Science, Department of Biological Sciences, Kuwait University, Safat, Kuwait, 13060; Department of Bioinformatics and Genetics, Swedish Museum of Natural History, Box 50007, SE-104 05 Stockholm, Sweden; Genome Center and Department of Evolution and Ecology, University of California - Davis, Davis, CA 95616, USA; Department of Veterinary Medicine and Surgery, College of Veterinary Medicine, University of Missouri-Columbia, Columbia, MO, 65211

**Keywords:** recombination, rho, recombination hotspot, domestic cat, *Felis silvestris catus*

## Abstract

Recombination has essential roles in increasing genetic variability within a population and in ensuring successful meiotic events. The objective of this study is to (i) infer the population scaled recombination rate (*ρ*), and (ii) identify and characterize localities of increased recombination rate for the domestic cat, *Felis silvestris catus.* SNPs (*n* = 701) were genotyped in twenty-two cats of Eastern random bred origin. The SNPs covered ten different chromosomal regions (A1, A2, B3, C2, D1, D2, D4, E2, F2, X) with an average region size of 850 Kb and an average SNP density of 70 SNPs/region. The Bayesian method in the program *inferRho* was used to infer regional population recombination rates and hotspots localities. The regions exhibited variable population recombination rates and four decisive recombination hotspots were identified on cat chromosome A2, D1, and E2 regions. No correlation was detected between the GC content and the locality of recombination spots. The hotspots enclosed L2 LINE elements and MIR and tRNA-Lys SINE elements in agreement with hotspots found in other mammals.

## Introduction

Recombination is a major source of genetic variation within sexually reproducing organisms, and is necessary for the proper alignment and segregation of homologous chromosomes during meiosis. Variation is achieved via recombination when new combinations of parental alleles across loci are generated and transmitted to succeeding generations. Lack of recombination may cause failure of meiotic division, or formation of gametes with chromosome number abnormalities (aneuploidy), which are often detrimental.

Localized chromosomal regions with elevated recombination rates are referred to as recombination “hotspots” (Steinmetz *et al*, 1982). Known hotspots in mice and human are generally 1 - 2 kb regions of high recombination rates surrounded by regions of low recombination (Paigen and Petkov, 2010). In humans, recombination hotspots are distributed about every 200 Kb (McVean *et al*, 2004) and over 25 000 hotspots have been identified (Myers *et al*, 2005). The first human recombination hotspot was identified using sperm typing via PCR in a region that harbors GC-rich mini-satellite (MS32) as a molecular signature (Jeffreys *et al*, 1998).

Several genomic features exhibit correlations with recombination hotspots. The GC content has been found to be positively correlated with recombination hotspots in humans (Myers *et al*, 2005), dog (Axelsson *et al*, 2012), pig (Tortereau *et al*, 2012), and chicken (Groenen *et al*, 2009) but not in mice (Wu *et al*, 2010). Long terminal repeats (LTR), long interspersed elements (LINE), and short interspersed elements (SINE) were also observed to be positively correlated with the locality of recombination hot spots in human (Lee *et al*, 2011; Myers *et al*, 2005), mice (Wu *et al*, 2010) and pigs (Tortereau *et al*, 2012). DNA motifs (*cis*) have been identified to be associated with recombination hot spots both in humans (Myers *et al*, 2005; Myers *et al*, 2008; Zheng *et al*, 2010) and in other organisms (Comeron *et al*, 2012). In addition to the *cis* elements presented by the motifs, a *trans* element, PRDM9, was found to be a major determinant of recombination hotspots in human and mice (Baudat *et al*, 2010; Brunschwig *et al*, 2012) but not in dog and related wild relatives (Auton *et al*, 2013; Munoz-Fuentes *et al*, 2011). PRDM9 is thought to bind to a DNA motif via a zinc finger domain, altering the chromatin structure through methylation, and recruiting recombination molecular machinery (Myers *et al*, 2010).

Coarse-scale recombination in cats has previously been investigated through linkage map analyses using sparsely distributed microsatellite markers (Menotti-Raymond *et al*, 1999; Menotti-Raymond *et al*, 2009). The objective of this study is to investigate fine-scale recombination rates, and recombination hotspots, in the domestic cat using population-level data for dense SNPs in ten selected genomic regions on ten different chromosomes.

## Materials and methods

### Samples and genotypes

SNP genotype data of twenty-two cats of Eastern random bred origin were obtained as previously described in (Alhaddad *et al*, 2013). The data were generated using a custom illumina GoldenGate array that represent ten different cat chromosomal regions (A1, A2, B3, C2, D1, D2, D4, E2, F2, X) and were composed of 1536 markers nearly equally distributed across the ten regions. Markers’ positions were updated to their location in the 6.2 cat genome assembly (http://genome.ucsc.edu/). To ensure successful inference, three criteria were used to filter the SNPs: (i) SNPs mapped to a single chromosomal location on the most recent genome assembly of cat with 100% sequence match, (ii) SNPs exhibiting genotype rate of ≥ 80%, and (iii) SNPs possessing a minor allele frequency of ≥ 0.1. The final dataset, included in this study, is composed of 701 markers distributed over the ten regions. The genomic locations of the regions and related summaries are shown in Table 1 (see also Figure 1 and Supplementary Information 2).

**Figure 1.**
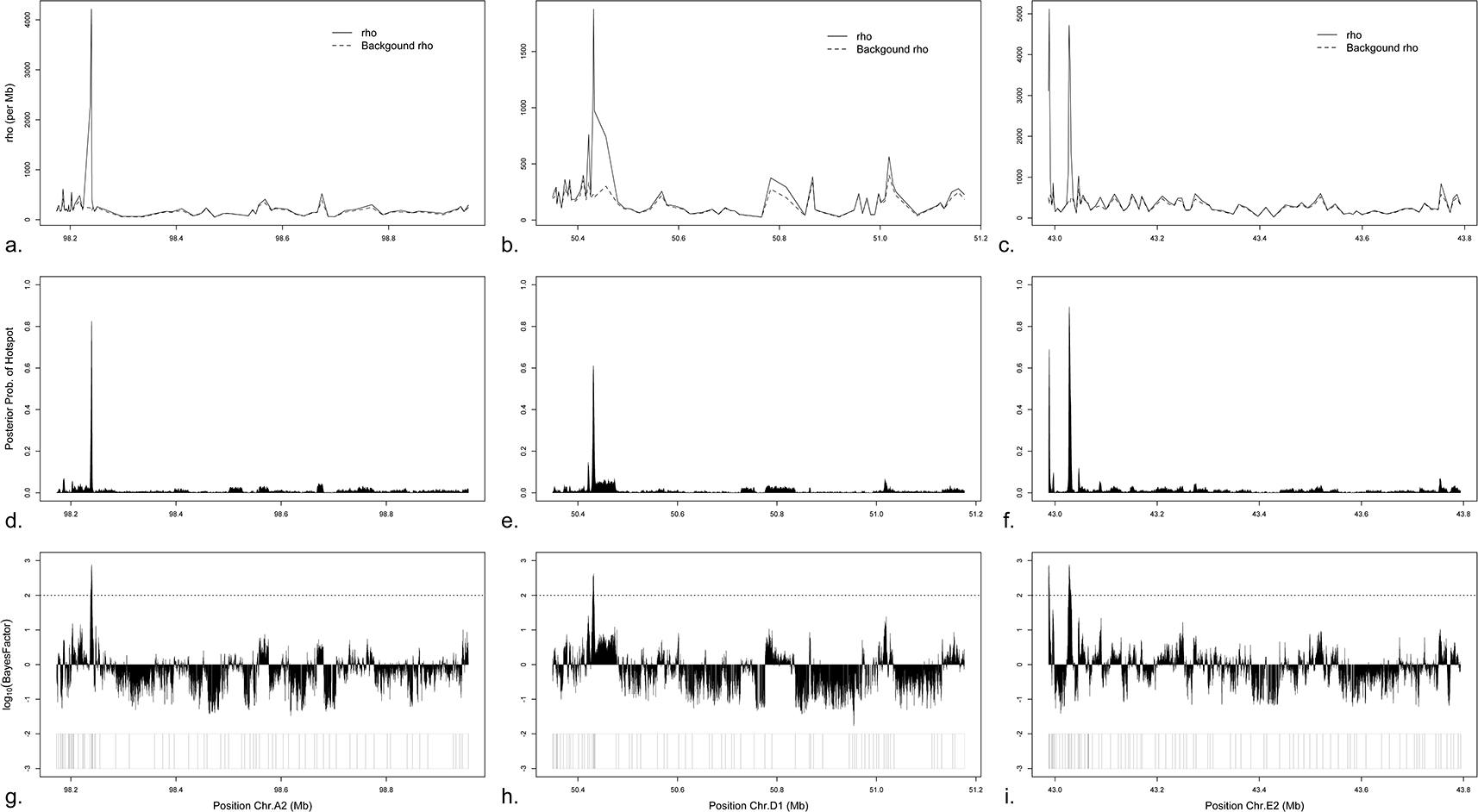
Recombination overview of chromosomes A2, D1, and E2 regions. (a-c) Posterior recombination rates (population size scaled) across chromosomes A2, D1, and E2 regions, respectively. Solid line shows the whole recombination rates, while the dashed line shows the background recombination rates. (d-f) Posterior probability of hotspots along chromosomes A2, D1, and E2 regions, respectively. (g-i) Bayes factor of hotspots for chromosomes A2, D1, and E2 regions, respectively. Horizontal dotted line corresponds to Bayes factor of 100 in a log_10_ scale. Position and distribution of SNPs included in the recombination analysis are at the bottom.

Three SNPs residing in a recombination hotspot on chromosome E2 region (see results) were chosen for genotype validation via sequencing. Primers were designed to flank the SNPs (Supplementary Information 2) and PCR was performed using DNA Engine Gradient Cycler (MJ Research, GMI, Ramsey, MN) using the following conditions. For each reaction, 2μl of DNA was used in a 1.5mM magnesium concentration with 1μM primers in a total reaction volume of 20μl. The annealing temperature of all primers was 62°C. The PCR protocol was as follows: initial denaturation at 94°C for 5 min followed by 40 cycles of 94°Cx45 sec, 62°Cx20 sec, 72°Cx30 sec, and a final extension at 72°C for 20 min. The PCR products were purified with ExoSap (USB, Cleveland, OH) per the manufacture’s recommendations and directly sequenced using the BigDye terminator Sequencing Kit v3.1 (Applied Biosystems, Foster City, CA) as previously implemented (Bighignoli *et al*, 2007). Sequences were verified and aligned using the software sequencer version 4.10 (Gene Codes Corp., Ann Arbor, MI).

## Recombination inference

The program *inferRho* uses the coalescent with recombination model (Hudson, 1991; Kingman, 1982a; Kingman, 1982b) in a full Bayesian framework to infer recombination rates and hot spots along the chromosome regions (Wang and Rannala, 2008; Wang and Rannala, 2009). The evolutionary relationship of the SNPs is represented by the ancestral recombination graph (ARG), which is an unobserved random variable that is integrated over using Markov Chain Monte Carlo (MCMC). In the variable recombination rate model, the crossing-over rate is *ρ_i_* = 4*N_e_c_i_,* where *N_e_* is the effective population size, *c_i_* is the recombination rate per generation in cM/Mb between marker *i* and *i* +1 (*i* = 1,…, *k* – 1, and *k* is the total number of SNPs). The total recombination rate (*c_i_*) consists of the background crossing-over rate (following a prior gamma distribution) and recombination hotspots (arising according to a Markov process) (Wang and Rannala, 2008; Wang and Rannala, 2009).

To make it computationally tractable, each chromosomal region was divided into blocks, with 20 SNPs per block (e.g., chromosome A1 region with 53 SNPs was divided into 3 blocks, 20 SNPs in the first and second block and 13 SNPs in the third block). Blocks from each chromosomal region are assumed to share the same population size parameter (*θ*) but have an independent ARG for each block. Preliminary runs were executed to determine the appropriate length and thinning interval of the MCMC chain for each chromosomal dataset. The last half of the MCMC samples (1000 samples) was used to estimate the recombination rates (*ρ_i_*) and to plot them with respect to the SNP marker positions. The distance-weighted mean recombination rate for each chromosomal region was calculated using the following equation:

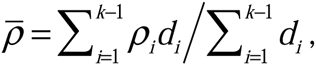

in which *d_i_* is the distance between marker *i* and *i*+1, and *k* is the total number of markers in the region.

## Designation of recombination hot spots

We calculated the Bayes factors (Kass and Raftery, 1995) to locate the positions of hotspots and represent relative odds of a hotspot being present. Each chromosomal region was divided into bins (200 bp per bin) for analysis of posterior samples to estimate the probability of having hotspot (*p_j_*) for each bin. The corresponding probability of hotspot for each bin (*q_j_*) from the prior was obtained by running the program without data (e.g., constant likelihood) but maintaining the same sample size and marker positions. The Bayes factor (*BF*) is defined as the ratio of the posterior and prior odds:

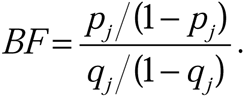

The total number of bins depends on the length of the region, thus the range of *j* is variable among the ten chromosome regions. Recombination hotspots are defined to have at least two consecutive bins with *BF* ≥ 100 (cf. section 3.2 in Kass & Raftery (1995)).

## GC content and genomic elements analyses

The GC content was calculated for the sequences of each bin using the function *CG.content* of package *APE* in R (Paradis *et al*, 2004).

Variation and repeat elements within each of the ten chromosomal regions were downloaded from UCSC genome browser using *RepeatMasker* for the v6.2 cat genome assembly. Elements were analyzed separately for bins with *BF* < 100 and *BF* ≥ 100 (hotspot bins). The elements within neutral bins (*BF* < 100) were to provide the general overview of the elements within each chromosomal region.

## Results

### Samples and genotypes

The dataset is composed of 701 SNPs on ten different cat chromosomal regions with an average of 70 SNPs/region. The chromosome E2 region harbors the highest number of SNPs (*n* = 90) while chromosome X region has the lowest (*n* = 37). SNPs are distributed across the regions with an average distance between SNPs of 12 Kb and a range of 145 bp – 651 Kb (Table 1). Four SNPs were selected for genotype verification using direct sequencing. The sequencing results showed were concordant to the genotypes obtained previously using the genotyping platform.

## Recombination rates and hotspots

The ten regions on chromosomes: A1, A2, B3, C2, D1, D2, D4, E2, F2, and X, were analyzed using *inferRho.* The lengths of the MCMC chains were determined as: 600 000 iterations for A1, C2; 800 000 iterations for A2, D1, D4, F2; 1 000 000 iterations for B3, D2, E2; and 500 000 iterations for X. Two thousand posterior samples for each chromosomal dataset were obtained after thinning, while the first half were discarded as burn-in. Using a parallel computing approach, each run could be accomplished within one to two weeks on a cluster with 2 Opteron 270 (2.0 GHz) processors per node.

Population recombination rate (*ρ*) is plotted for each region (Figure 1 a-c, Supplementary Information 2a). The mean recombination rate (*ρ*) across all regions is 200 per Mb. The E2 region exhibits the highest mean rate (309 /Mb) whereas X region has the lowest (77 /Mb) (Table 1). The difference between the background recombination rate and whole recombination rate is indicative of recombination spots. This difference is most noticeable in regions of chromosomes A2, D1, and E2 (Figure 1 a-c).

The posterior probability of hotspot was calculated for bins of size 200 bp in each region (Figure 1 d-f, Supplementary Information 2b). Chromosomes A2, D1, and E2 show distinctly high posterior probabilities (> 0.6) in four localized areas (Figure 1 d-f). The posterior probabilities for the other seven chromosome regions (A1, B3, C2, D2, D4, F2, and X) are less than 0.2, indicating little support for elevation of recombination rate in these areas.

The Bayes factors (Figure 1 g-i, Supplementary Information 2c) are consistent in pattern with the posterior probabilities (Figure 1 d-f, Supplementary Information 2b) and recombination rates (Figure 1 a-c, Supplementary Information 2a). Approximately 99% (*n* = 42,531) of bins were classified as “neutral” (*BF* < 100) across all regions examined. The hot spots were found in only three chromosomal regions (A2, D1, and E2) and represented ~ 0.13% (*n* = 57) of all bins studied. Summaries of the numbers and distribution of bins within each chromosomal region are provided in Table 1. The hotspots had size of 3 Kb in A2, 1.8 Kb in D1, and 1.8 Kb for the first and 4.6 Kb for the second in E2. The distance between the two hotspots in E2 was ~37.4 Kb (Table 2).

## GC content analysis

The log_10_ of the Bayes factor was plotted as a function of the GC content in Supplementary Information 3. Pearson’s correlation test revealed a positive correlation between GC and log_10_(Bayes factor) (cor = 0.077, *p* < 0.0001) but was not suggestive of strong correlation. Moreover, no significant differences in the mean GC content of each class of bins were observed (t-test, *p* = 0.05) (Supplementary Information 3). The mean GC contents of the hotspots are shown in Table 2.

## Repeat elements analysis

The four hotspot regions contained 22 repeat and variation elements (Table 2, Figure 2a). SINE elements constitute the highest proportion (40%) of the elements present in the hotspots followed by LINE elements (27%). Within LINE elements, L2 elements were present in three of the four hotspot regions. MIR family elements were present in all hotspot regions and tRNA-Lys family elements are present in three of the four elements. Low complexity, long terminal repeats, simple repeats, and DNA elements were inconsistently present across the hotspot regions.

**Figure 2.**
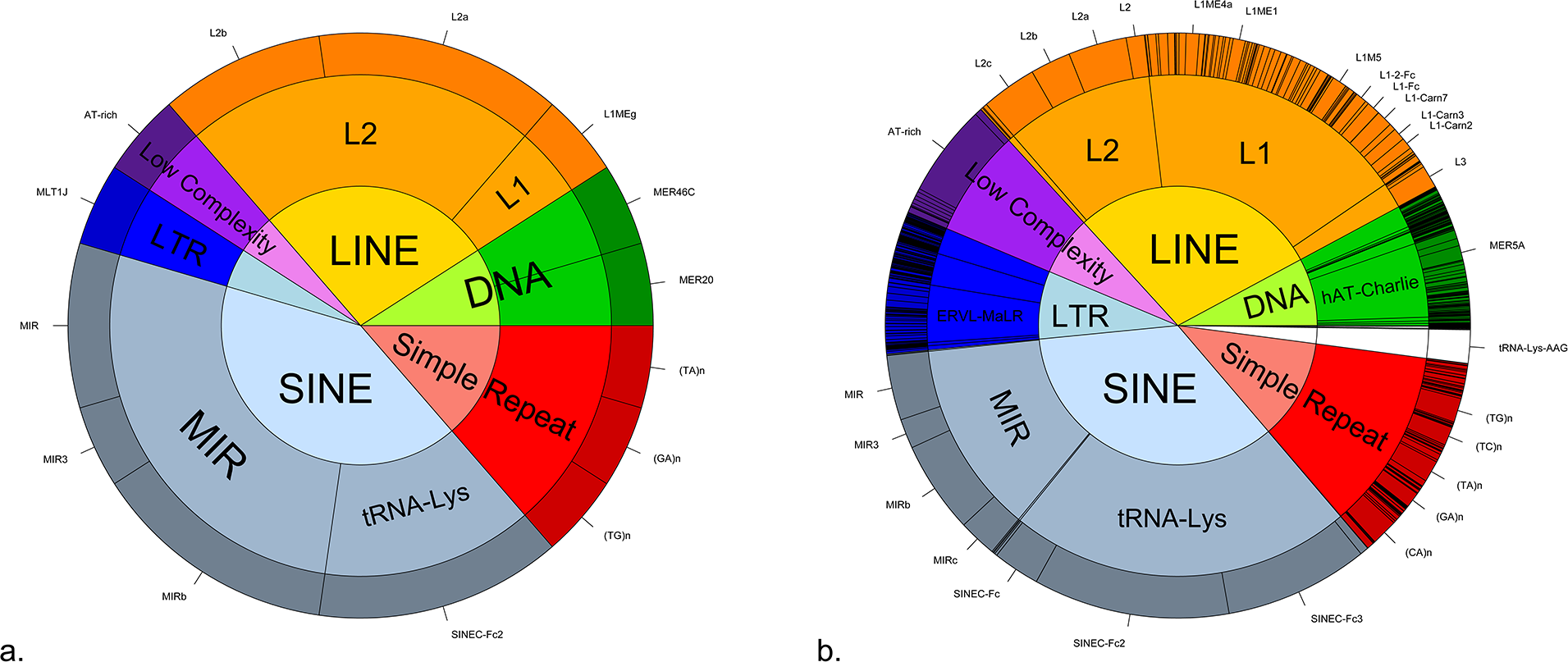
Overview of variation and repeat elements in the cat chromosomal regions studied. a) Variation and repeat element (*n* = 22) within four hotspots. All elements within the hotspots are shown at the tips of the outer circle. b) Variation and repeat element (*n* = 16,798) within “neutral” bins. Elements present >100 within the warm-spots are shown at the tips of the outer circle. The inner circle represents the elements’ classes, middle circle represents elements’ families, and outer circle represent the individual elements.

Variation and repeat elements within neutral regions where investigated to get a general view of the general distribution of repeat elements (Figure 2b). In neutral regions, the SINE elements represent the highest proportion of elements, 34 %. The tRNA-Lys SINE elements constitute 22% and MIR SINE elements represent 12%. The second highest repeat elements was the LINE elements, 29%, where L1 elements represent ~17% and L2 elements represent 9.5% of all elements in the neutral regions.

## Discussion

Advances in population genetic theory and technology have made it possible to estimate recombination rates directly from genotype data on population samples, overcoming the limitations of sperm typing or using large extended families (Hellenthal and Stephens, 2006). The strategy of the population genetics based approach is to use information on the number of recombination events that have occurred in the history of the population, which can be detected by modeling the patterns of genetic variation expected to be present in randomly selected individuals.

The model accounts for coalescent and recombination events in an Ancestral Recombination Graph (ARG). Markers will have coalescent trees that are likely to vary across the genome. In theory, all markers in a chromosome are correlated by an ARG. However, the size of the ARG might grow much faster than linear with the increasing number of SNPs, making it computational intractable. In practice, the data are usually partitioned into blocks. Blocks from the same chromosomal region are assumed to share the same population size parameter (*θ*) but have an independent ARG for each block. For fast computation, some methods use an approximate likelihood instead of the full likelihood calculation, for example, the composite-likelihood method implemented in *LDhat* (McVean *et al*, 2002) and PAC-likelihood method implemented in *PHASE* (Stephens and Donnelly, 2003). Approximate-likelihood methods may be feasible to apply to large genomic regions, but may lack power to detect a moderate or low rate of recombination. Full-likelihood method in *inferRho* uses all of the information in the data and should therefore provide more accurate estimates (Wang and Rannala, 2008; Wang and Rannala, 2009).

The analysis of recombination hotspots in cats, presented here, constitutes the first application of the program *inferRho* to non-human data and the first analysis of fine-scale recombination rates in cats. The population recombination rate was found to be variable between the regions analyzed and, as expected, the mean recombination rate of X chromosome regions was lower than that of any autosomal regions. This variation in recombination rates and the notable reduced rate outside of the pseudoautosomal region of the X chromosome are in agreement with observations of recombination in human (Myers *et al*, 2005) and dog (Axelsson *et al*, 2012). The latter result is expected due to that fact that recombination outside of the pseudoautosomal region occurs only in females.

Four decisive hot spots were identified on three chromosomes: A2, D1 and E2. The localities of the hotspots are in agreement with the localities of increased posterior probabilities and the general topography of recombination rates as expected. Acknowledging the limitation posed by the total size of the regions analyzed (total ~ 8.5 Mb) compared to the size of the genome and the lack of power to perform correlation and element enrichment analyses, the following observations have been made: (i) Unlike hotspots in humans (Myers *et al*, 2005), dog (Axelsson *et al*, 2012), pig (Tortereau *et al*, 2012), and chicken (Groenen *et al*, 2009), which exhibit positive correlation with GC content, the hotspots in cats, identified in this study, show no distinct positive correlation with GC content (this is similar to mice) (Wu *et al*, 2010). (ii) Genomic features in the four hot spots are consistent with those found in other mammals. Three of the decisive hotspots contained at least one L2 LINE element, which is consistent with features of hotspots in human and chimpanzee (Lee *et al*, 2011; Myers *et al*, 2010; Myers *et al*, 2008). Similarly, MIR and tRNA-Lys SINE elements were found in hotspots of cats, which is consistent with the positive correlation found for hotspots in human (Lee *et al*, 2011), and pig (Tortereau *et al*, 2012). (iii) The general similarity of the hotspot repeat elements to those of other mammals suggests similar recombination mechanism and probable involvement of PRDM9. (iv) This similarity suggests there may be evolutionary differences between cats and dogs, especially since recent studies found that PRDM is not functional in dog (Axelsson *et al*, 2012; Munoz-Fuentes *et al*, 2011) and that recombination hotspots in dogs are associated with CpG and promoter regions (Auton *et al*, 2013).

This study represents a glimpse of recombination hotspots in cats and only an initial step toward understanding recombination in cats. As cat resources developing, genome-wide analyses may be performed and more definitive conclusions will reach. Nonetheless, understanding the recombination landscape will shed light on the mechanism of recombination in cats compared to other species, the pattern of variation generated by recombination in cats, and should lead to better implementation of efficient disease mapping strategies in cats.

## Acknowledgments

We would like to thank Drs. Jeffery Ross-Ibarra, Robert A. Grahn, and Barbara Gandolfi for their comments and suggestions. This project has been funded in part previously by the National Center for Research Resources R24 RR016094 and is currently supported by the Office of Research Infrastructure Programs OD R24OD010928, the Winn Feline Foundation (W10-014, W11-041), the George and Phyllis Miller Feline Health Fund, and the Center for Companion Animal Health, School of Veterinary Medicine, University of California, Davis. HA is funded with a full scholarship by Kuwait University.

## Reference

Alhaddad H, Khan R, Grahn RA, Gandolfi B, Mullikin JC, Cole SA et al (2013). Extent of linkage disequilibrium in the domestic cat, Felis silvestris catus, and its breeds. Plos One 8(1): e53537.

Auton A, Rui Li Y, Kidd J, Oliveira K, Nadel J, Holloway JK et al (2013). Genetic recombination is targeted towards gene promoter regions in dogs. ArXiv e-prints **arXiv**:1305.6485.

Axelsson E, Webster MT, Ratnakumar A, Ponting CP, Lindblad-Toh K (2012). Death of PRDM9 coincides with stabilization of the recombination landscape in the dog genome. Genome Res 22(1): 51–63.

Baudat F, Buard J, Grey C, Fledel-Alon A, Ober C, Przeworski M et al (2010). PRDM9 is a major determinant of meiotic recombination hotspots in humans and mice. Science 327(5967): 836–840.

Bighignoli B, Niini T, Grahn RA, Pedersen NC, Millon LV, Polli M et al (2007). Cytidine monophospho-N-acetylneuraminic acid hydroxylase (CMAH) mutations associated with the domestic cat AB blood group. BMC Genet 8: 27.

Brunschwig H, Levi L, Ben-David E, Williams RW, Yakir B, Shifman S (2012). Fine-scale Map of Recombination Rates and Hotspots in the Mouse Genome. Genetics 112.141036

Comeron JM, Ratnappan R, Bailin S (2012). The many landscapes of recombination in Drosophila melanogaster. PLoS Genet 8(10): e1002905.

Groenen MA, Wahlberg P, Foglio M, Cheng HH, Megens HJ, Crooijmans RP et al (2009). A high-density SNP-based linkage map of the chicken genome reveals sequence features correlated with recombination rate. Genome Res 19(3): 510–519.

Hellenthal G, Stephens M (2006). Insights into recombination from population genetic variation. Curr Opin Genet Dev 16(6): 565–572.

Hudson RR (1991). Gene genealogies and the coalescent process. Oxford Survey in Evolutionary Biology 7: 1–44.

Jeffreys AJ, Murray J, Neumann R (1998). High-resolution mapping of crossovers in human sperm defines a minisatellite-associated recombination hotspot. Mol Cell 2(2): 267–273.

Kass RE, Raftery AE (1995). Bayes Factors. Journal of the American Statistical Association 90(430): 773–795.

Kingman JFC (1982a). The coalescent. Stochastic Processes and their Applications 13(3): 235–248.

Kingman JFC (1982b). On the Genealogy of Large Populations. Journal of Applied Probability 19: 27–43.

Lee YS, Chao A, Chen CH, Chou T, Wang SY, Wang TH (2011). Analysis of human meiotic recombination events with a parent-sibling tracing approach. BMC Genomics 12: 434.

McVean G, Awadalla P, Fearnhead P (2002). A coalescent-based method for detecting and estimating recombination from gene sequences. Genetics 160(3): 1231–1241.

McVean GA, Myers SR, Hunt S, Deloukas P, Bentley DR, Donnelly P (2004). The fine-scale structure of recombination rate variation in the human genome. Science 304(5670): 581–584.

Menotti-Raymond M, David VA, Lyons LA, Schaffer AA, Tomlin JF, Hutton MK et al (1999). A genetic linkage map of microsatellites in the domestic cat (Felis catus). Genomics 57(1): 9–23.

Menotti-Raymond M, David VA, Schaffer AA, Tomlin JF, Eizirik E, Phillip C et al (2009). An autosomal genetic linkage map of the domestic cat, Felis silvestris catus. Genomics 93(4): 305–313.

Munoz-Fuentes V, Di Rienzo A, Vila C (2011). Prdm9, a major determinant of meiotic recombination hotspots, is not functional in dogs and their wild relatives, wolves and coyotes. Plos One 6(11): e25498.

Myers S, Bottolo L, Freeman C, McVean G, Donnelly P (2005). A fine-scale map of recombination rates and hotspots across the human genome. Science 310(5746): 321–324.

Myers S, Bowden R, Tumian A, Bontrop RE, Freeman C, MacFie TS et al (2010). Drive against hotspot motifs in primates implicates the PRDM9 gene in meiotic recombination. Science 327(5967): 876–879.

Myers S, Freeman C, Auton A, Donnelly P, McVean G (2008). A common sequence motif associated with recombination hot spots and genome instability in humans. Nat Genet 40(9): 1124–1129.

Paigen K, Petkov P (2010). Mammalian recombination hot spots: properties, control and evolution. Nat Rev Genet 11(3): 221–233.

Paradis E, Claude J, Strimmer K (2004). APE: Analyses of Phylogenetics and Evolution in R language. Bioinformatics 20(2): 289–290.

Steinmetz M, Minard K, Horvath S, McNicholas J, Srelinger J, Wake C et al (1982). A molecular map of the immune response region from the major histocompatibility complex of the mouse. Nature 300(5887): 35–42.

Stephens M, Donnelly P (2003). A comparison of bayesian methods for haplotype reconstruction from population genotype data. Am J Hum Genet 73(5): 1162–1169.

Tortereau F, Servin B, Frantz L, Megens HJ, Milan D, Rohrer G et al (2012). A high density recombination map of the pig reveals a correlation between sex-specific recombination and GC content. BMC Genomics 13: 586.

Wang Y, Rannala B (2008). Bayesian inference of fine-scale recombination rates using population genomic data. Philos Trans R Soc Lond B Biol Sci 363(1512): 3921–3930.

Wang Y, Rannala B (2009). Population genomic inference of recombination rates and hotspots. Proc Natl Acad Sci U S A 106(15): 6215–6219.

Wu ZK, Getun IV, Bois PR (2010). Anatomy of mouse recombination hot spots. Nucleic Acids Res 38(7): 2346–2354.

Zheng J, Khil PP, Camerini-Otero RD, Przytycka TM (2010). Detecting sequence polymorphisms associated with meiotic recombination hotspots in the human genome. Genome Biol 11(10): R103.

